# Determination of chemical partitioning in vehicles using a skin surrogate for dermal absorption model refinement

**DOI:** 10.1101/2025.05.29.656843

**Authors:** Ruth Pendlington, David Sanders, David Sheffield, Stephen Glavin, Hugh Barlow, Hequn Li

**Affiliations:** Safety, Environmental and Regulatory Sciences (SERS) group, Unilever, Colworth Science Park, Sharnbrook MK44 1LQ, United Kingdom

## Abstract

In nonclinical safety assessments of cosmetic ingredients applied topically, it is essential to investigate the rate and extent of dermal penetration to reach the systemic circulation. The initial step in skin absorption involves the partitioning of an ingredient from the formulation into the upper layer of the stratum corneum and this plays a key role in determining the delivery of an ingredient. Currently, there is a lack of reliable methods to accurately predict this important parameter. To address this, we measured the partitioning of three chemicals from seven different vehicles into a surrogate PDMS membrane; these measurements were then used to parameterize a dermal absorption model. The evaluation of receptor fluid kinetics and skin absorption percentages, comparing measured and predicted data, demonstrated that incorporating the vehicle partitioning parameter derived from the PDMS system improved the accuracy of model predictions for the three chemicals in most vehicles.

## INTRODUCTION

Accurately determining the absorption of drugs or ingredients into the skin is crucial for assessing systemic exposure in topically applied products. In vitro skin absorption studies, following established guidelines, provide reliable estimates of absorption and aid in safety risk assessment. Ex vivo human tissue is advantageous in cosmetics safety assessment as it provides human-relevant data and can be used with physiologically-based kinetic (PBK) modelling to predict the fate of absorbed cosmetic ingredients. (Bernauer et al. 2023). Additionally, ex vivo skin penetration data can be used with physiologically-based kinetic (PBK) modelling to predict the fate of absorbed cosmetic ingredients (Li et al. 2022). However, in vitro studies are both financial and human resource-intensive, and using ex vivo human skin raises additional consent and ethical concerns, given the use of human material. Radiolabelling ingredients enables sensitive measurements of skin absorption, but custom synthesis and limited stability of radiochemicals pose challenges. Furthermore, data generated from one study cannot be universally applied to different products containing the same ingredient.

The reason for this is that formulation can have a great impact on how an ingredient will interact with the skin (Otto et al. 2009a; Pendlington et al. 2008). The first step of skin absorption is partitioning of the ingredient out of the vehicle in which it has been applied to the skin and into the stratum corneum. The physicochemical properties of the ingredient, and those of the formulation in which it finds itself, will determine whether the ingredient partitions easily into the lipid rich stratum corneum or not. A cosmetic ingredient may be used in a wide range of formulations e.g. skin cream, shampoo, underarm deodorant etc which have very different physicochemical properties. Even within a cosmetic type, formulations may differ widely, e.g. skin creams and lotions may be water-in-oil, oil-in-water or even water-in-oil-in-water, so even within a single product type, skin absorption of a specific ingredient may be markedly different depending on the formulation properties (Supe and Takudage 2021). Running an *ex vivo* skin absorption experiment for every product type or every variation within a product type is clearly not practical.

If the first step in the process of skin absorption (i.e. partitioning of an ingredient out of the vehicle of the formulation and into the upper layer of the stratum corneum) could be predicted or measured, the necessity to perform skin absorption studies for an ingredient of interest from different formulations may no longer be necessary. To our best knowledge, Quantitative Structure Activity Relationship (QSAR) approaches are not available for predicting such vehicle effect related properties (Grégoire et al. 2021). A number of groups have published on using isolated stratum corneum to measure partition coefficients of various chemicals, (Hansen et al. 2008), (Ellison et al. 2020) and others, however preparation of isolated stratum although not difficult, does take time, human resource and requires the precious commodity of human ex vivo skin being available, so we decided to investigate a more “off the shelf” approach. Polydimethylsiloxane membranes (silastic membranes) have been used by various groups in place of skin to assess the skin absorption of materials (Chilcott et al. 2005), (Van der Merwe and Riviere 2005). By adding a chemical into a system involving the immersion of a PDMS membrane in a specific vehicle, it allows for the measurement of the chemical’s partitioning between the PDMS and the vehicle. This measurement of ingredient partitioning using PDMS could enable the ranking of formulation efficacy in delivering actives to the skin. Moreover, these measurements can be utilized to derive parameter values on vehicle partitioning for a dermal absorption model.

In this study, we aimed to address the following objectives: we 1) investigated the influence of different vehicles on skin absorption by conducting an ex vivo skin penetration study of three chemicals (caffeine, coumarin, and 4HR) across eight different vehicles. Specifically, we examined the distributions of the chemicals in various skin layers and their absorption profiles in the receptor fluid; 2) we described and implemented a method that utilizes a readily available non-biological PDMS membrane to measure the partition coefficients of the three chemicals between PDMS and seven of the vehicles, this enabled us to calculate an input parameter related to vehicle partitioning for refining a dermal absorption model using GastroPlus; 3) then we compared the refined predictions obtained using the PDMS-derived partition coefficients with measured ex vivo human skin absorption data. This comparison allowed us to assess the practical utility of the PDMS-derived parameter for predicting the partitioning of chemicals into vehicles and its impact on skin absorption predictions.

## MATERIALS & METHODS

### Materials

#### Chemicals

4-Hexyl [U-^14^C]resorcinol and [3-^14^C]coumarin were supplied by Pharmaron, Cardiff, UK; [1-Methyl-^14^C]caffeine was supplied by American Radiolabeled Chemicals Inc., St. Louis, MO, USA. Analytical grade absolute ethanol and HPLC grade Ultrapure water were supplied by VWR. Polydimethylsiloxane (PDMS) sheets (Silicone Elastomer) 1mm thick were supplied by Goodfellow, Cambridge, UK. A low water (6%) containing face serum base was supplied by Unilever Trumbull, CT, USA, a Shampoo was supplied by Unilever Port Sunlight, UK (PS Shampoo, used for the skin absorption study and initial PDMS measurements). Caffeine, coumarin, 4-hexylresorcinol and olive oil were supplied by Sigma-Aldrich, Gillingham, UK. The Body Lotion (Vaseline Intensive Care Lotion Aloe Soothe) and the second Shampoo (used for the later PDMS measurements, Alberto Balsam Coconut & Lychee) were purchased from a local supermarket.

#### Skin samples

Fifteen samples of full-thickness human skin (fourteen abdominal and one abdomen/arm) were obtained from fifteen donors (fourteen female and one male) aged 36 to 76 years old patients who gave informed consent for their skin to be taken for scientific purposes prior to undergoing routine surgery. Three samples were obtained from Bio-repository, NHS Greater Glasgow & Clyde (REC reference 16/WS/0207), eight samples from Tissue Solutions and five samples from NHS Lothian. Samples that arrived frozen were stored in a freezer at -20°C. Skin samples that arrived on ice had subcutaneous fat and connective tissue removed using a scalpel, the skin was washed under cold running tap water, dried with paper towels, cut into pieces, wrapped in aluminium foil or vacuum-packed and stored in a freezer at -20^°^C. Human skin samples were removed from -20°C storage and allowed to thaw at ambient temperature. Split-thickness membranes were prepared by pinning the full-thickness skin, stratum corneum uppermost, onto a raised cork board and cutting with an electric dermatome (Zimmer®) at a setting equivalent to 200-400 µm depth. The thickness of the membranes was measured using a micrometer. Membranes were then wrapped in foil, placed into a self-sealing bag and stored in a freezer at -20°C for a maximum period of two months.

#### Skin absorption measurements

Skin samples were mounted in flow-through diffusion cells (0.64 cm^2^ dose application area), which were set in a heated steel manifold to give a skin surface temperature of 32°C. Receptor fluid (phosphate buffered saline containing 5% (v/v) human serum, amphotericin B (10 mL of a 250 μg/mL solution per litre of receptor fluid), streptomycin (ca 0.1 mg/mL) and penicillin (ca 100 units/mL)) was pumped through the cells at 1.5 ml/h for a 15-min equilibration period before dosing. The solubility of each test item in the receptor fluid indicated that infinite sink conditions would be maintained throughout the experiment. Barrier integrity was assessed by electrical resistance using a Tinsley Databridge (Model 6401); any skin sample exhibiting a resistance <10.9 kΩ was excluded from subsequent absorption measurements.

See Table 1 for a summary of amounts dosed and application times. The test items in all test preparations were at 0.5% w/w (5mg/g), the shampoo formulation was then diluted ten-fold with water prior to application. Test preparations were applied evenly over the surface of the exposed skin of 10 split-thickness samples of human skin using a positive displacement pipette set to deliver 5 mg/cm^2^, apart from the diluted Shampoo Base formulation which was applied at 50 mg/cm^2^. The donor chambers of the cells were occluded with traps containing carbon filters, apart from those applied with the Shampoo Base formulation, which were left unoccluded. Six representative weighed aliquots of each test preparation were dispensed into vials at the time of dosing, mixed with methanol: scintillation fluid and analysed by liquid scintillation counting. The radiochemical purity of the radiolabelled test items was determined post dose using radio-HPLC.

**Table 1.**
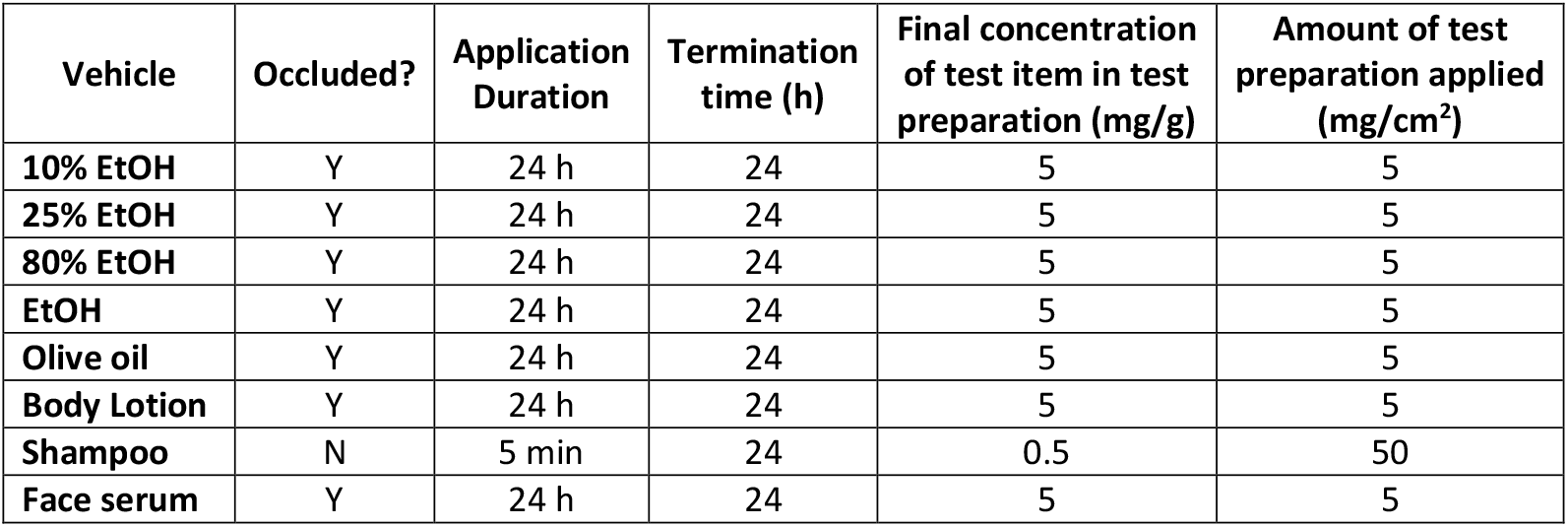
Summary of dosing conditions.

| Vehicle | Occluded? | Application Duration | Termination time (h) | Final concentration of test item in test preparation (mg/g) | Amount of test preparation applied (mg/cm<sup>2</sup>) |
| --- | --- | --- | --- | --- | --- |
| 10% EtOH | Y | 24 h | 24 | 5 | 5 |
| 25% EtOH | Y | 24 h | 24 | 5 | 5 |
| 80% EtOH | Y | 24 h | 24 | 5 | 5 |
| EtOH | Y | 24 h | 24 | 5 | 5 |
| Olive oil | Y | 24 h | 24 | 5 | 5 |
| Body Lotion | Y | 24 h | 24 | 5 | 5 |
| Shampoo | N | 5 min | 24 | 0.5 | 50 |
| Face serum | Y | 24 h | 24 | 5 | 5 |

Application time for the shampoo formulation was 5 min, after which the formulation was removed with a tissue swab. The surface of the skin was washed with water (5 mL in 0.5 mL aliquots) and dried with a tissue swab; this process was then repeated. For every other dose group, application time was until the termination time point. Absorption of the test item was assessed by collection of receptor fluid in hourly fractions from t = 0 until the termination time point.

The termination time point for every dose group was 24 h (n = 5 cells). The underside of the skin was rinsed with receptor fluid and the rinsing retained. Each diffusion cell was then disconnected from the receptor fluid inlet pump lines and removed from the heated manifold. The traps and filters were separated and extracted with solvent; in all cases the extraction solvent for caffeine and coumarin was ethanol, for 4-hexylresorcinol it was methanol.

For the Shampoo Base formulation dose group, the skin surface was washed again with water as detailed above, for all other formulations the skin was rinsed with 5 mL of 2% (v/v) commercial soap solution in 0.5 mL aliquots, dried with a tissue swab and the process repeated. Pipette tips used to aspirate the rinsings were retained.

The donor and receptor chambers were dismantled, the skin removed, then the chambers were each extracted with solvent. The stratum corneum was removed with 20 successive tape strips (D-Squame® tape discs); each tape strip was analysed individually. The dosed area of skin was separated from the unexposed skin, wrapped in cling film and a 200 g weight heated to 65°C in a water bath placed onto the epidermal surface for 90 s. The epidermis was then peeled away from the dermis using a scalpel. The epidermis, dermis and unexposed skin samples were placed into separate vials containing Solvable® and heated at 60°C to aid solubilisation; when fully dissolved, stannous chloride solution (0.2 g/mL in ethanol; 150 μL) was added to the skin samples. All retained samples were analysed by liquid scintillation counting.

#### Partition coefficient measurements

Partition coefficients were measured in seven of the eight formulations used in the skin absorption study; these experiments were run at a later date than the skin absorption study and unfortunately fresh samples of the same face serum were unavailable at that time.

Aliquots of stock solutions of 4-HR in absolute ethanol or caffeine in 50% v:v chloroform/absolute ethanol were transferred into 7ml glass vials and dried down under nitrogen; 5ml vehicle was then added. For coumarin, which is volatile, 100µl aliquots of stock solution in absolute ethanol were added to each vial and 4.9ml vehicle added. All vials were shaken in an incubator set at 20^°^C using a pulse mode shaker, 500rpm/0.5sec pulse. After 72 hours, in vials where the test item had completely dissolved, the vials were removed for ^14^C assay. Remaining vials were shaken for a further 32 hours before standing for approximately 16 hours for the undissolved solids to settle (4-HR), or for a further 48 hours followed by centrifugation at 4000rpm for 10 minutes in a bench centrifuge to pellet the undissolved solids (coumarin and caffeine). The final concentration (target 0.5 %, saturated if 5mg/g vehicle not achievable) and homogeneity of the test item in the loading vehicle was measured by liquid scintillation counting (three weighed aliquots taken); homogeneous preparations were defined as having a coefficient of variation ≤ 5 %. The Body Lotion samples were not homogeneous with respect to ^14^C, so further attempts were made to homogenise the samples, including centrifugation and using a handheld Omni THQ homogeniser. The Omni THQ homogeniser (15,000rpm in 2 × 10 minute bursts) followed by centrifugation at 4000rpm for 90 minutes greatly improved the homogeneity assay of the body lotion samples, and although not every sample met the acceptance criteria, it was decided to include them in the study.

Discs 1cm in diameter were cut from a 1mm thick sheet of PDMS and solvent cleaned by rolling in HPLC grade methanol for approximately 8 hours. The discs were air-dried overnight at ambient. Five PDMS discs were added to small lengths of stainless-steel wire.

One wire was placed into each vial, so that all discs were immersed in the vehicle. Where test item hadn’t fully dissolved, care was taken to leave the solids undisturbed. The vials were shaken at 55rpm in an incubator set at 20^°^C. In the first experiments, after 3, 6, 24, 48, 120 and 168 hours, one vial from each vehicle group was taken for the assay to determine how long it took for equilibration to occur (First Study). In subsequent experiments (First and Second Study) a single incubation time of 96 hours was used. The discs were removed individually from the wire, dipped briefly in ultrapure water, wiped with lint-free cloth and weighed. Each disc was then allowed to stand for approximately 24 hours in 10ml of Pico Fluor Plus scintillation cocktail before liquid scintillation counting (LSC). Where solids remained, these were disturbed during removal of the discs and therefore these vials were centrifuged at 4000rpm for 40 minutes to re-pellet the solids. After removing the discs, each loading vehicle (or the supernatant where solids had been pelleted) was analysed by LSC. Aliquots (1ml, n=5) of each blank vehicle were weighed to enable the concentration of test item to be calculated as µg/g. The mean PDMS/solvent partition coefficient for each time point was calculated as:

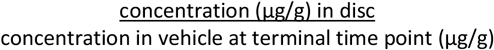

Following in-house development of the method (First Study and Second Study), a further study was outsourced to a Contract Research Organisation (CRO) to determine the transferability of the method (Outsourced Study). The conditions employed in that study were those employed in the Second Study, the only difference being an Ultra Turax T 10 hand-held homogeniser was used in place of the Omni THQ homogeniser.

### Assessment of variability in the data sets

To assess the variability within the data sets, the results from the Outsourced Study were analysed. There are several sources of variation (see Figure 1) that need to be considered when estimating the partition coefficient (Disc concentration / Vehicle concentration).

**Figure 1.**
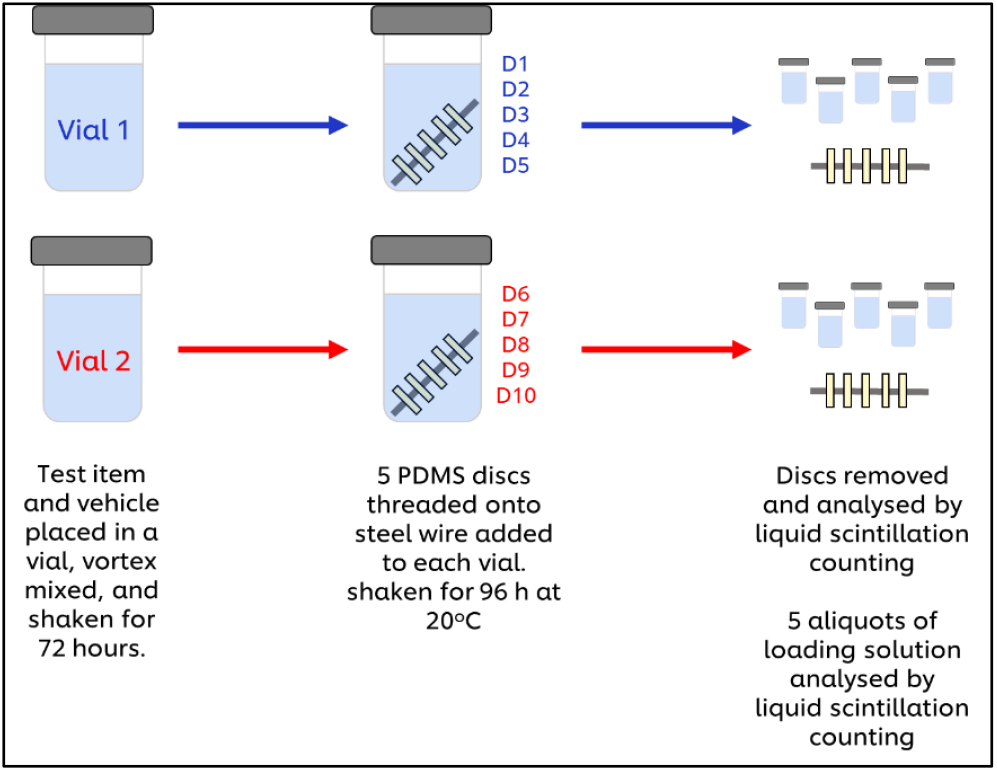

- between disc variability.
- between vehicle aliquot variability.
- between vial variability.

To account for all these sources of variation, a model was set up with terms for disc concentration, vehicle concentration and vial effect. Also, because partition coefficient is a ratio of disc and vehicle concentrations, the equation needs to be transformed from a multiplicative model to an additive one by taking logs of the concentration, then back transform the difference between disc and vehicle estimates to get the partition coefficient.

**Equation 1:** Partition Coefficient = Disc Concentration / Vehicle Concentration

**Equation 2:** Log_10_(Partition Coefficient) = Log_10_(Disc Concentration) – Log_10_(Vehicle Concentration)

**Equation 3:** Partition Coefficient = 10^(Log_10_(Disc Concentration) – Log_10_(Vehicle Concentration)) The model was fit using the Mixed procedure in SAS 9.4 (The SAS Institute).

**Equation 4:** Log_10_(Concentration) = Type(i) + Vial(j) Where:

*i* = Disc or Vehicle.

*j* = Vial 1 or Vial 2.

### Dermal absorption modelling

Dermal absorption modelling of the three chemicals was conducted using GastroPlus 9.8 (Simulation Plus, Lancaster, CA, USA). The GastroPlus dermal module (TCAT) in GastroPlus is a complex mechanistic model of dermal absorption, which can simulate a variety of transdermal dosage forms, including liquid formulations (solutions, lotions, suspensions) and semi-solid formations (gels, creams, lotions, pastes). The transdermal drug delivery model represents the skin as a collection of the following compartments (Figure 2): stratum corneum, viable epidermis, dermis, subcutaneous tissue, sebum, hair lipid, and hair core, to account for the drug concentration gradient within each compartment due to drug diffusion under non-steady-state conditions.

**Figure 2.**
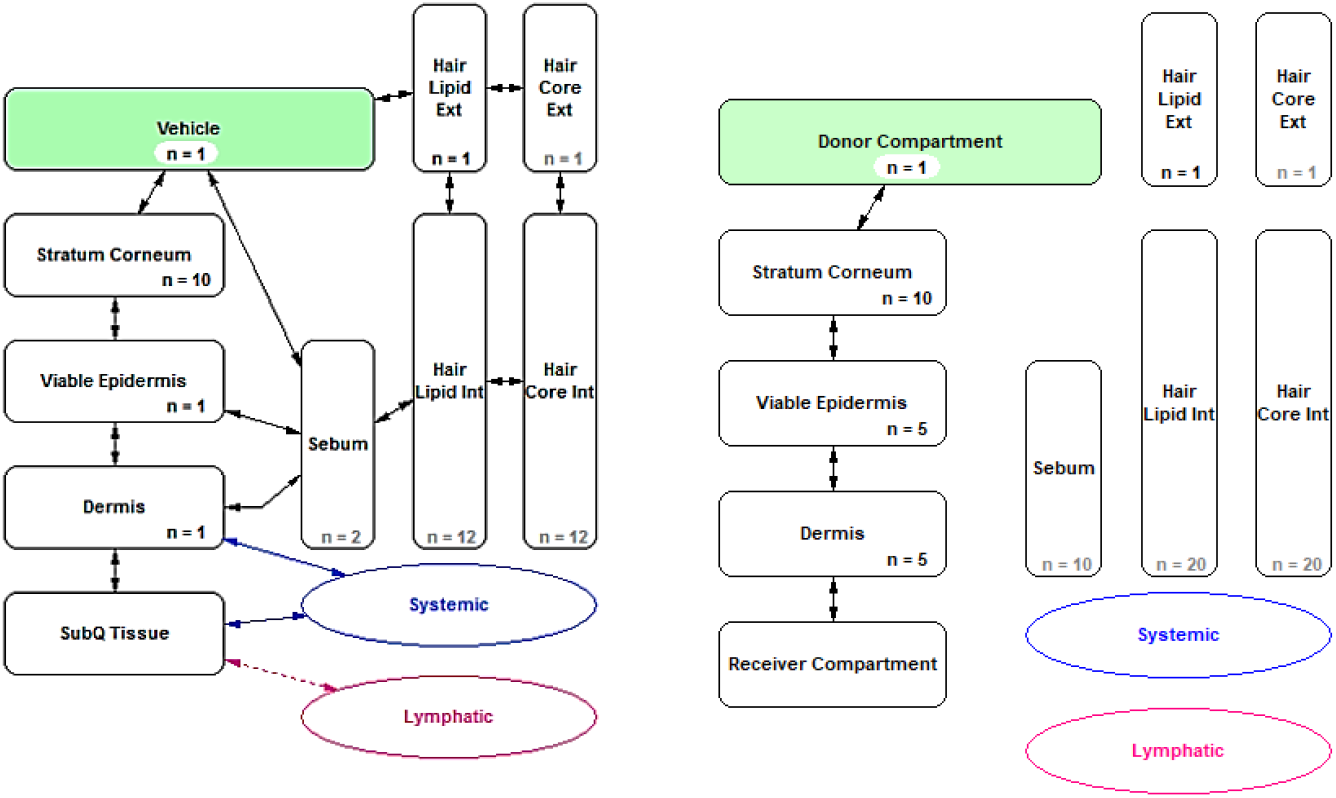
(Left) schematic diagram of the TCAT module in GastroPlus showing how the different compartments are connected to one another and the rest of the body and (right) the compartments involved for modelling in vitro skin penetration.

To examine the usefulness of integrating PDMS-derived data in the dermal absorption model on formulation effect, a model on dermal absorption was built incorporating a list of input parameters including the chemical/skin interaction parameters like diffusion and partitioning coefficients of the chemical in various skin layers, i.e. the stratum corneum, epidermis, dermis, vehicle related parameters, such as vehicle-water partition coefficient, evaporation, solubility in solvent etc. and dosing scenario related parameters.

To this end, Human-In Vitro Abdomen was chosen as the dermal physiology in TCAT and donor compartment, stratum corneum, viable epidermis, dermis and receiver compartment were included to mimic the in vitro skin penetration experiment. Parameters like dose, volume, applied surface area, application time and whether the skin is covered were input into the model to mimic the ex vivo skin penetration scenario. The chemical/skin interaction parameters, i.e. the partition coefficient between a skin layer and water, as well as diffusivities in skin layers, i.e. stratum corneum, epidermis and dermis, were obtained by fitting to the ex vivo skin penetration data (kinetic in receptor fluid and distribution in skin layers data) from 10% ethanol solution. Then the fitted parameters were used as inputs for other vehicle solutions. For the vehicle-water partition coefficient, it was either parameterised as the default value (1 in TCAT) or derived from PDMS data (as follows) for comparison,

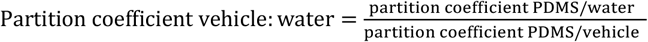

Apart from the vehicle partitioning parameters, another important formulation effect is chemical or vehicle loss due to evaporation. As the diffusion cells were occluded (except for the shampoo formulation) significant chemical loss was not expected. However, from the skin absorption study results, in some of the ethanol water vehicle conditions, certain loss of coumarin was seen. Therefore, the loss of chemical was accounted for by setting the initial dose multiplied by the mass balance percentage for the situations where mass balance was below 85%.

Evaporation of vehicles or the solubilities of chemicals in different vehicles were not measured, so the related parameters were either predicted or estimated based on assumptions. The first assumption made was when occluded, water/oil content loss can be neglected, so that for the body lotion and olive oil conditions no evaporation is modelled. However, the ethanol content loss needs to be modelled. This was based on our previous measured data of the fate of ethanol in skin absorption experiment for 24 hours that only 40% was recovered, even though the skin was covered (occluded) (Pendlington et al. 2001). Therefore, for the ethanol containing vehicles, it was assumed that 60% of the ethanol content will evaporate, so the residual volume fraction is 0.94, 0.85, 0.52 and 0.4 for 10%, 25%, 80% and 100% ethanol as the vehicle. For the shampoo condition (unoccluded), it was assumed the main volume loss is due to water evaporation, so that the residual volume is 0.5 for this condition. To model the evaporation of ethanol (for the 4 ethanol containing vehicles) or water (for shampoo vehicle) over time, the evaporation rates were predicted using the Nielsen method in the TCAT module. The input parameters of Solvent vapor pressure were obtained from https://chem.libretexts.org/ and solvent diffusivity in air were obtained from reported literature values (Smith et al. 2017).

Solubility of the chemicals in the vehicle is an influential factor too, as different absorption profiles can result from infinite, finite, and saturated dosing conditions. For the 10%, 25% and 80% ethanol vehicles, as evaporation of ethanol to the headspace of the occluded condition is predicted to be fast (time for 60% of the ethanol content to evaporate is 0.7 hours (Water -Thermal Diffusivity vs. Temperature and Pressure (engineeringtoolbox.com)), the main solubility is driven by water. For the 100% ethanol and olive oil conditions, the solubility is driven by ethanol or oil for them being the sole solvent. And for the body lotion and Shampoo conditions, it is assumed the solubility in the vehicle is similar to that of water. Detailed information on the values used for modelling the vehicle evaporation and solubility in the solvent are shown in tables 2 and 3. The simulated dermal absorption profile, i.e. distribution in the skin layers and flux in receptor fluid, was then compared against the ex vivo skin penetration data for evaluation.

**Table 2.** Prediction on volume loss of vehicle due to solvent evaporation to headspace (occluded) or air (unoccluded) and the solubility in the residual volume of the vehicle.

| Vehicle | Occluded | Volatile | Volume loss driving solvent | Solvent Vapor pressure (Torr) | Solvent AirDiffusivity (m<sup>2</sup>/s) | AirKinematic Viscosity (m<sup>2</sup>/s) | Evaporation rate Model | AirVelocity (m/s) | Solvent evaporation rate (ml/s) | Residual Volume Fraction | Solubility driving solvent | Solvent MW (g/mol) | Solvent Density | Solvent pH |
| --- | --- | --- | --- | --- | --- | --- | --- | --- | --- | --- | --- | --- | --- | --- |
| 10% EtOH | Y | Y | Ethanol | 40 | 1.5E-5 | 1.6E-5 | Nielsen | 1.0E-4 | 2.5E-2 | 0.94 | water | 18 | 1 | 7 |
| 25% EtOH | Y | Y | Ethanol | 40 | 1.5E-5 | 1.6E-5 | Nielsen | 1.0E-4 | 2.5E-2 | 0.85 | water | 18 | 1 | 7 |
| 80% EtOH | Y | Y | Ethanol | 40 | 1.5E-5 | 1.6E-5 | Nielsen | 1.0E-4 | 2.5E-2 | 0.52 | water | 18 | 1 | 7 |
| EtOH | Y | Y | Ethanol | 40 | 1.5E-5 | 1.6E-5 | Nielsen | 1.0E-4 | 2.5E-2 | 0.4 | ethanol | 46 | 0.7 | 7 |
| Olive oil | Y | N | n.a. | n.a. | n.a. | n.a. | n.a. | n.a. | n.a. | 1 | oil | 1500 | 0.92 | 7 |
| Body lotion | Y | N | n.a. | n.a. | n.a. | n.a. | n.a. | n.a. | n.a. | 1 | water | 18 | 1 | 7 |
| Shampoo | N | Y | Water | 17 | 1.4E-7 | 1.6E-5 | Nielsen | 0.44 | 7.6E-2 | 0.5 | water | 18 | 1 | 7 |

**Table 3.**
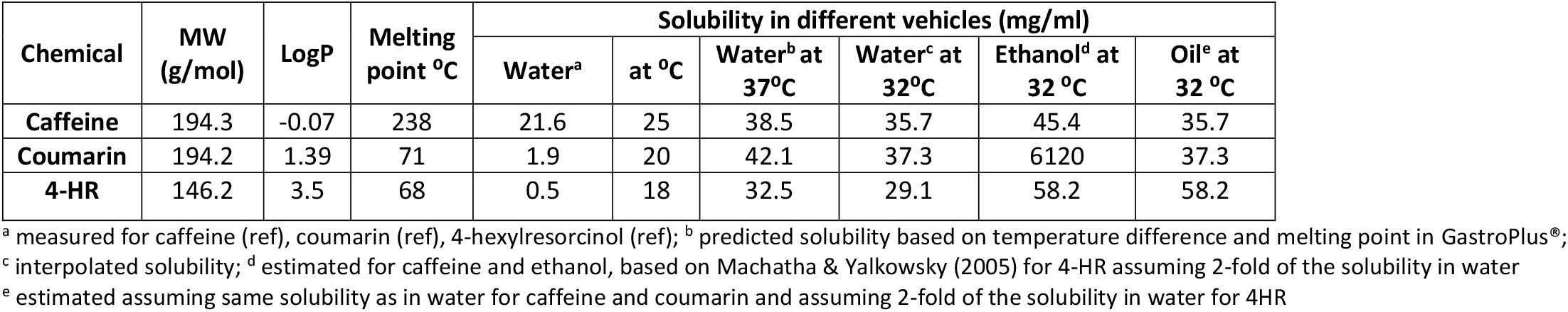
Table physico-chemical properties of the three chemicals.

## RESULTS

### Partition coefficient measurements

Equilibration of all three test items was rapid (graphs available in supplementary information appendix 1). The PDMS/solvent partition coefficients for caffeine were all below 0.1. For Coumarin, the more aqueous the solvent the greater the partition coefficient was. 4-Hexylresorcinol showed a distinct one to two orders of magnitude difference between the partition coefficient for the two most aqueous vehicles (water and 10% aqueous ethanol) and the remaining vehicles. In general, the data produced in the three separate studies were comparable (Table 4).

**Table 4.** PDMS/Solvent partition ratios measured in three studies.

| Test Item | Solvent | PDMS/Solvent<br>partition ratio<br>First Study | PDMS/Solvent<br>partition ratio<br>Second Study | PDMS/Solvent<br>partition ratio<br>Outsourced Study |
| --- | --- | --- | --- | --- |
| Caffeine | Water | 0.026** | ND | 0.075 |
| Caffeine | 10% Ethanol | ND | 0.079 | 0.039 |
| Caffeine | 25% Ethanol | 0.009** | 0.040 | 0.021 |
| Caffeine | 80% Ethanol | ND | 0.009 | 0.008 |
| Caffeine | Ethanol | 0.063** | 0.060 | 0.067 |
| Caffeine | Olive Oil | ND | 0.385 | 0.355 |
| Caffeine | 10% shampoo* | ND | 0.099 | 0.077 |
| Caffeine | Body lotion | ND | 0.037 | 0.040 |
| Coumarin | Water | 1.908**<br>1.887 | ND | ND |
| Coumarin | 10% Ethanol | ND | 1.628 | 1.540 |
| Coumarin | 25% Ethanol | 0.609** | 0.723 | 0.694 |
| Coumarin | 80% Ethanol | ND | 0.054 | 0.054 |
| Coumarin | Ethanol | 0.089** | 0.091 | 0.088 |
| Coumarin | Olive Oil | 0.124 | 0.199 | 0.182 |
| Coumarin | 10% shampoo* | ND | 1.898 | 1.970 |
| Coumarin | Body lotion | 0.902 | 1.112 | 0.961 |
| 4-HR | Water | 3.674**<br>1.885 | ND | ND |
| 4-HR | 10% Ethanol | ND | 2.665 | ND |
| 4-HR | 25% Ethanol | 0.212** | 0.359 | ND |
| 4-HR | 80% Ethanol | ND | 0.007 | ND |
| 4-HR | Ethanol | 0.022** | 0.024 | ND |
| 4-HR | Olive Oil | 0.019 | 0.018 | ND |
| 4-HR | 10% Shampoo* | ND | 0.085 | ND |
| 4-HR | Body lotion | 0.025 | ND | ND |

\*First Study and Second Study – PS shampoo; Outsourced Study – Supermarket shampoo
\*\* 168 h rather than 96 h incubation; ND Not done

### Assessment of variability in the data set (Outsourced study)

#### Vials

Graphical results can be found in Supplementary data S1. In most cases there was no difference between vial 1 and vial 2, only Oil with caffeine and 10% ethanol, ethanol, shampoo and body lotion with coumarin showed a significant vial effect, the difference was due to experimental variation.

#### Partition Coefficients

Supplementary data S2 Table 1 (caffeine) and Table 2 (coumarin) show the partition coefficient estimates and their 95% confidence limits. They are graphically represented in Figure 3 (caffeine) and Figure 4 (coumarin) below. The within material variance was relatively small compared to the between material variance, shown by the small confidence intervals in the figures.

**Figure 3:**
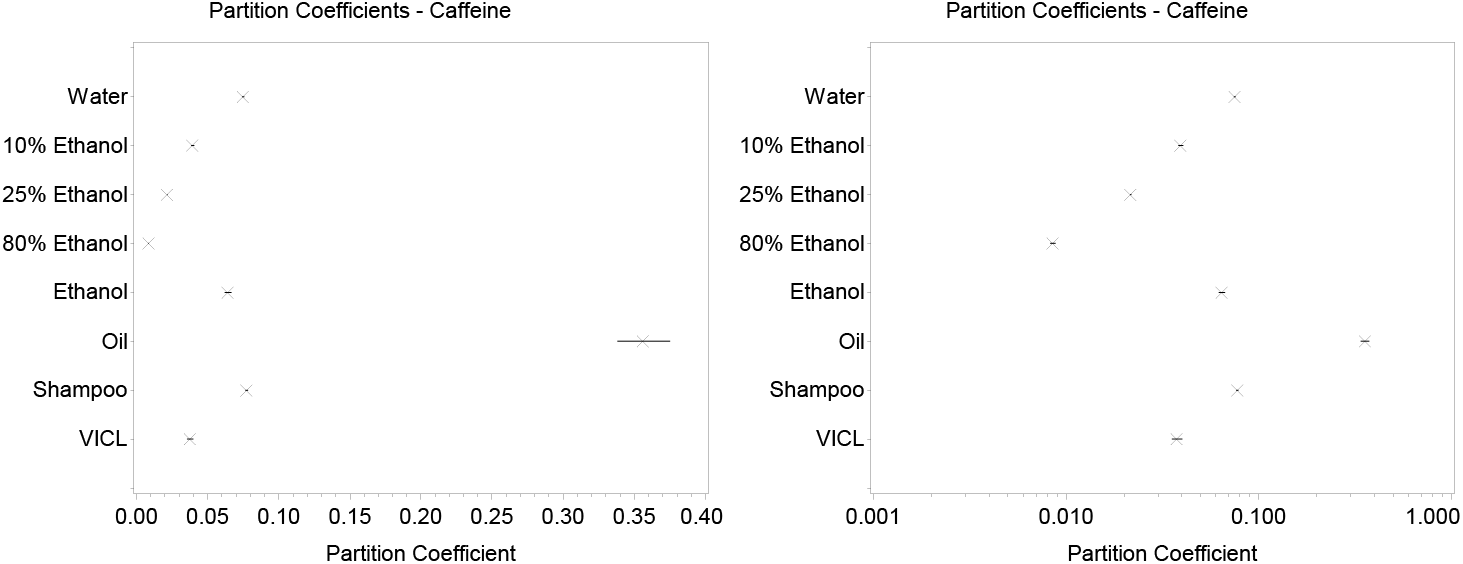
Partition coefficients for caffeine with 95% confidence interval (left: ordinary scale, right: log scale)

**Figure 4:**
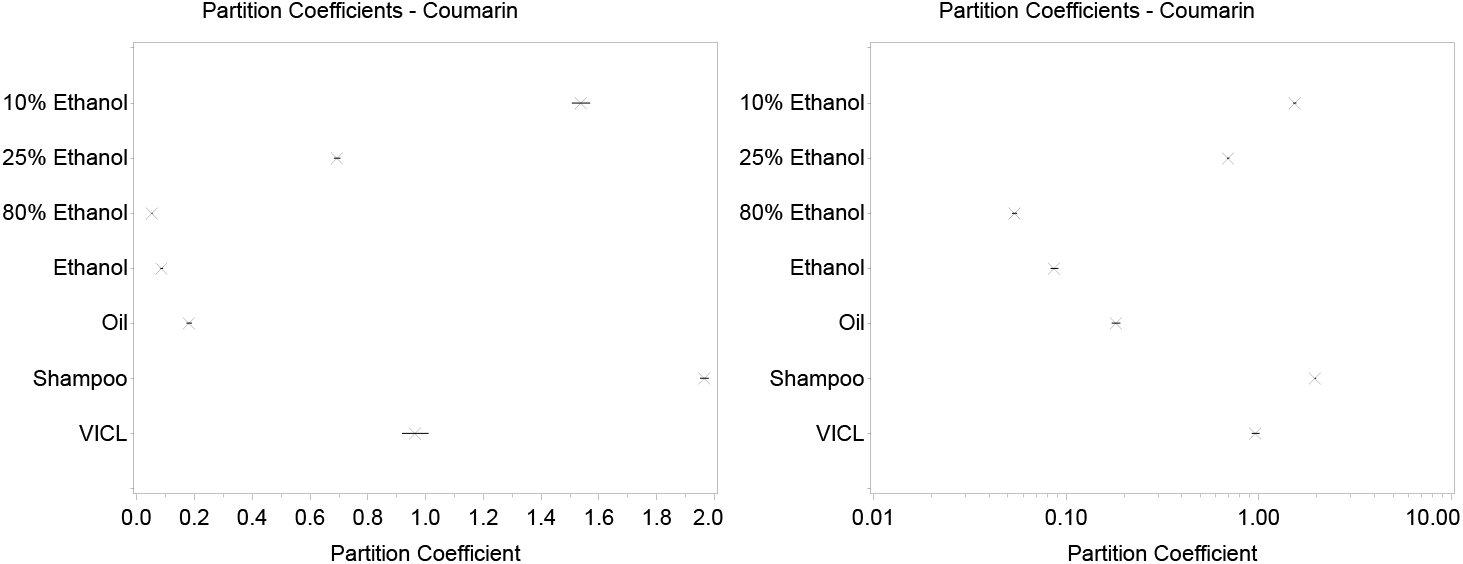
Partition coefficients for coumarin with 95% confidence interval (left: ordinary scale, right: log scale)

Oil was the only vehicle with caffeine with a 95% confidence interval larger than 0.01; it was also by far the largest partition coefficient for caffeine. For both caffeine and coumarin, as the proportion of ethanol increased, the partition coefficient decreased up to 80% ethanol, after which it increased for 100% ethanol, slightly for coumarin and considerably for caffeine, the latter increasing almost to the same value as its partition coefficient for water. Olive oil gave the highest partition value for caffeine, while shampoo had the highest value for coumarin.

### Skin penetration data

The distribution within the skin tissue at the 24-hour time point and the flux through the skin of each chemical from each formulation are presented in Figures 5-7.

**Figure 5.**
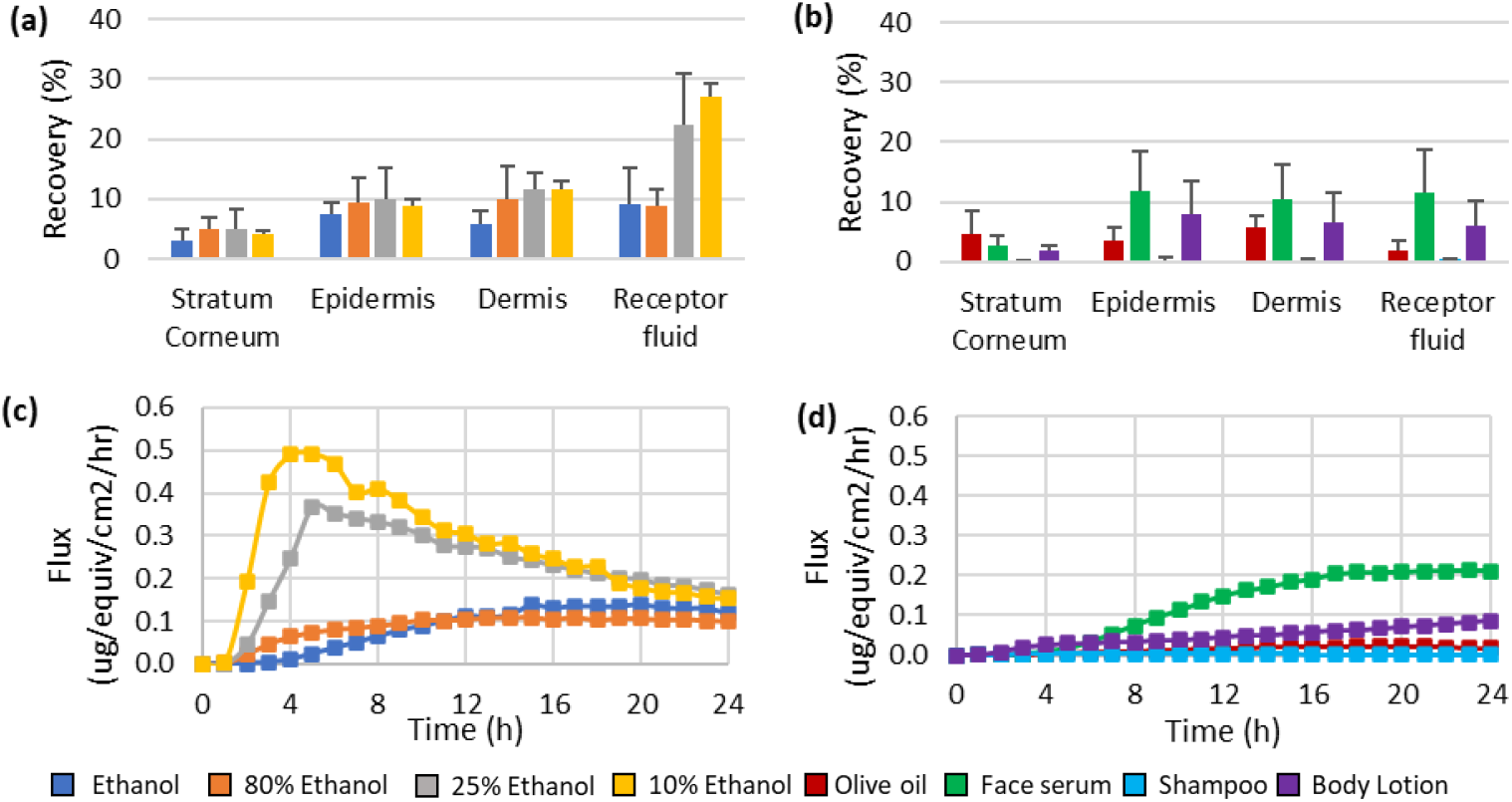
4-Hexylresorcinol skin absorption: (a) distribution at 24 h from application in water/ethanol vehicles; (b) distribution at 24 h from application in other vehicles; (c) flux through the skin into receptor fluid from water/ethanol vehicles; (d) flux through the skin into receptor fluid from other vehicles.

It can be seen that the extent of skin absorption differs across the different chemicals for each vehicle and across the different vehicles for each chemical. In general, coumarin seems to penetrate more than caffeine and 4HR do when the vehicle applied was the same across the three chemicals. When ethanol water mixtures were the vehicles, the percentage of ethanol present in the vehicle influenced the extent of skin penetration. For caffeine, a parabolic effect was observed with percentage absorption first increasing with an increase in the percentage of ethanol in the vehicle, then decreasing when applied in neat ethanol, whereas for coumarin and 4HR, the ethanol percentage seemed to have an inverse correlation with the chemicals’ absorption i.e. when the ethanol fraction went up, the amount absorbed decreased. When olive oil was used as the vehicle, the absorption for all three chemicals was lower than the ethanol-water vehicles, and the biggest impact was shown on 4HR. The Body Lotion showed to have different effects on the absorption depending on the chemical used; the absorption for caffeine and coumarin both went up when compared with most of the ethanol-water or oil conditions, however, the absorption stayed quite low for 4HR. The last vehicle is shampoo that for all three chemicals the absorption was rather low due to removal of chemical through rinsing off at 5min post dosing (Figure 8).

One thing to note about the data is the inherent variability seen in these experiments; each dose group was only n = 5 diffusion cells (due to practical and economic constraints), with n = 1 cell per skin donor per dose group; for the purposes of generating data for safety risk assessment purposes, we use a minimum of four donors per dose group, three cells per donor (i.e. n = 12 per dose group) in an attempt to reduce this variability. Data from the latter type of experiment show that the variability can be intra-donor as well as inter-donor (unpublished observations).

The second thing to note is the shape of the flux curve can enable a greater understanding of how a test item is absorbed. For example, in Figure 6c it can be seen that the dermal delivery of caffeine from ethanolic solutions increases to a maximum rate between about three and five hours, then decreases again, whereas its absorption rate from olive oil (Figure 6d) reaches steady state from eight hours, whilst absorption from the body lotion continues to increase throughout the experiment, perhaps just attaining steady state in the last three or so hours. For Coumarin, meanwhile, Figure 7c, passage into and through the skin was rapid for all of the ethanol/water vehicles, with practically nothing remaining in the skin fractions by 24 hours; the same pattern is seen for Coumarin’s absorption from body lotion, but the absorption rate is much slower from the other three vehicles (Figure 7d).

**Figure 6.**
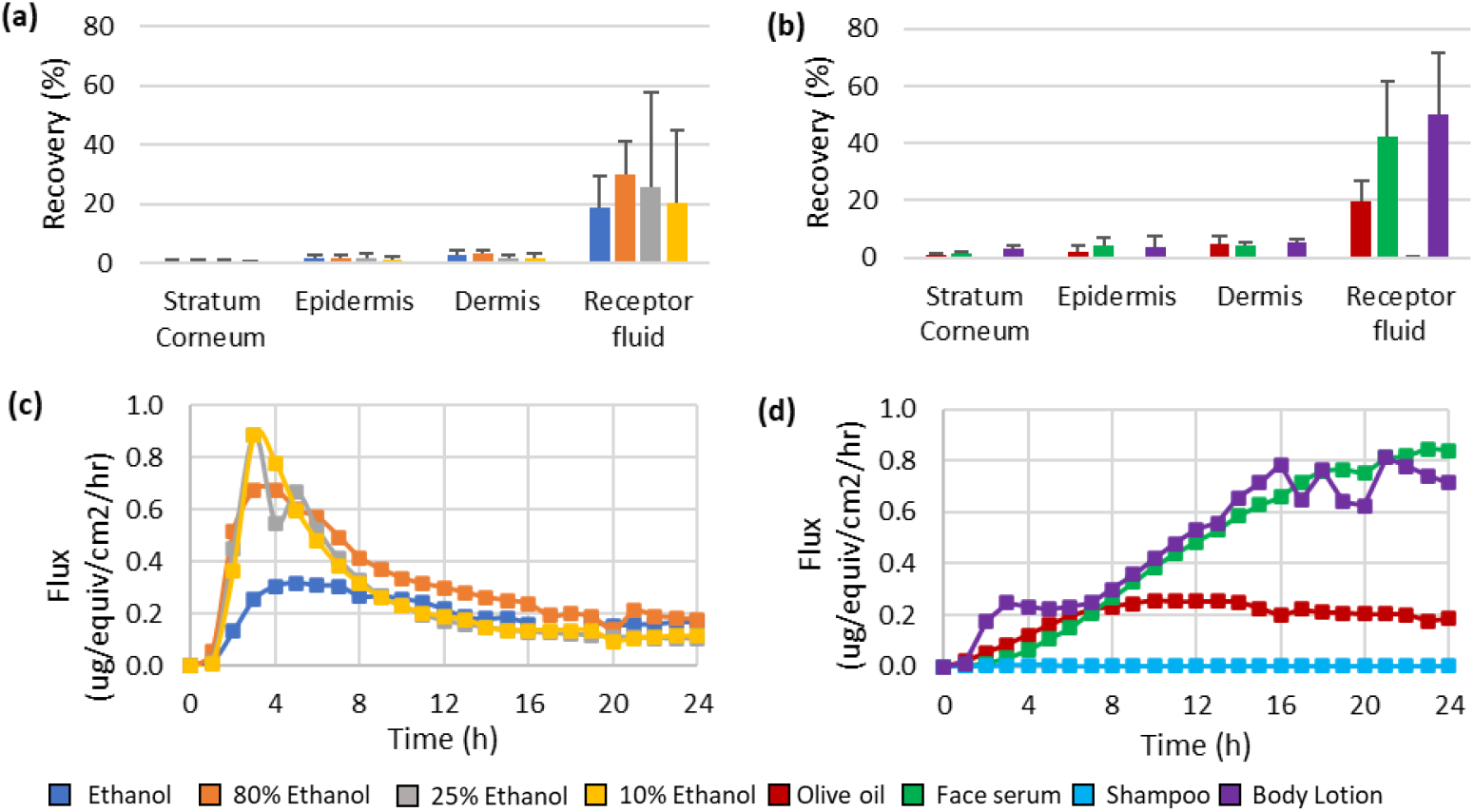
Caffeine skin absorption: (a) distribution at 24 h from application in water/ethanol vehicles; (b) distribution at 24 h from application in other vehicles; (c) flux through the skin into receptor fluid from water/ethanol vehicles; (d) flux through the skin into receptor fluid from other vehicles.

**Figure 7.**
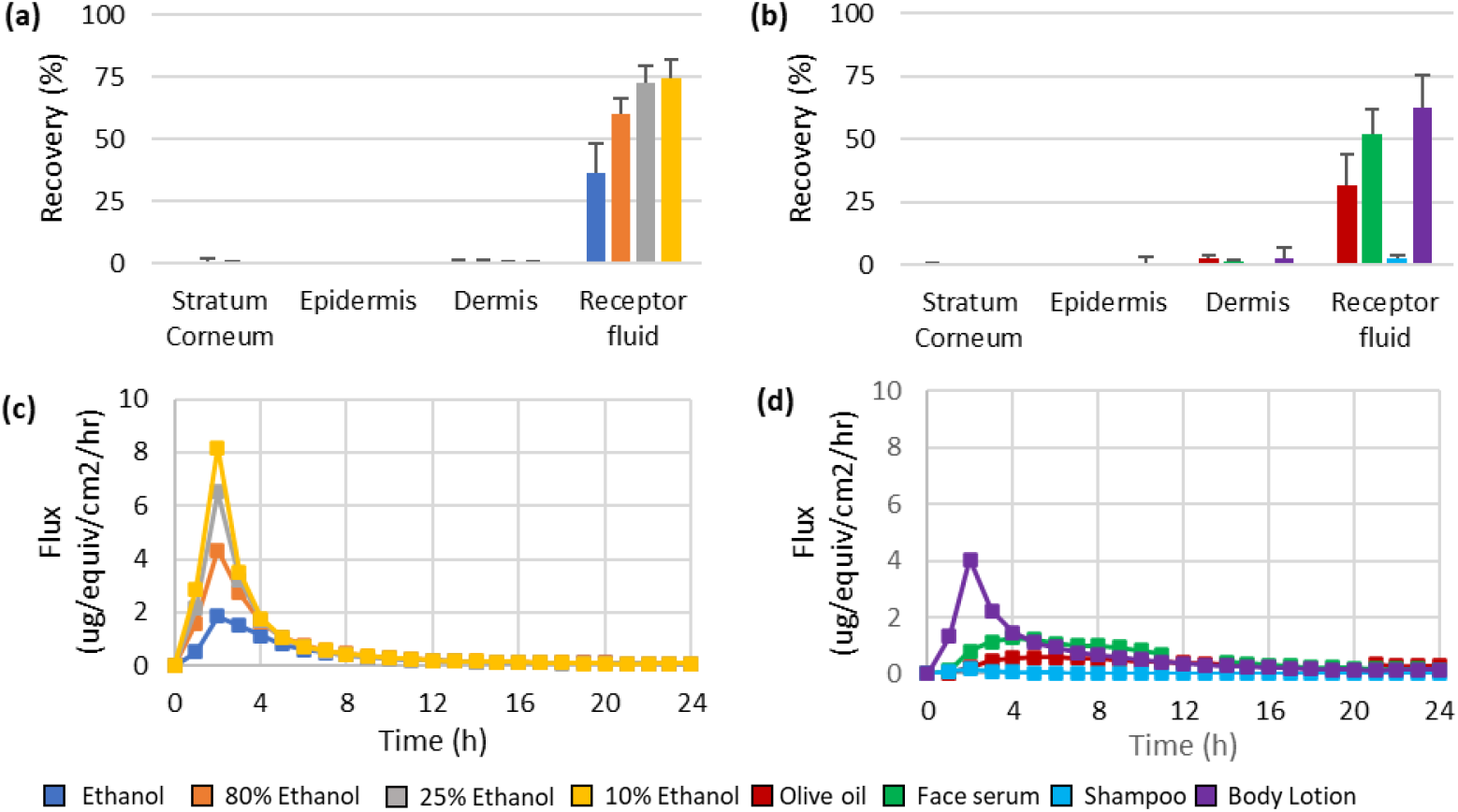
Coumarin skin absorption: (a) distribution at 24 h from application in water/ethanol vehicles; (b) distribution at 24 h from application in other vehicles; (c) flux through the skin into receptor fluid from water/ethanol vehicles; (d) flux through the skin into receptor fluid from other vehicles.

**Figure 8.**
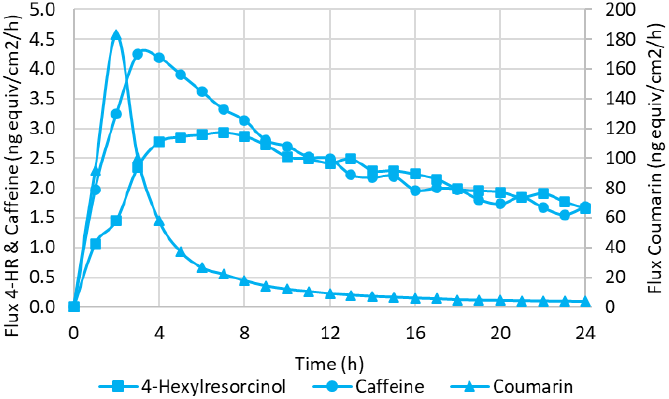
Flux of 4-Hexylresorcinol, caffeine and coumarin through the skin into receptor fluid from shampoo

Coumarin also had the poorest mass balance figures across the board of vehicles, with only the mass balance from the shampoo dose group fulfilling the 100±10% acceptable mass balance as set out in the OECD guideline 428 (OECD 2004) (Figure 9). Coumarin is volatile, hence a filter and trap were used to occlude the diffusions cells for all but the shampoo formulation, however very little material was recovered from the filter and trap for any dose group, indicating that the system was inefficient at trapping Coumarin that had volatilised off the surface of the skin. The recovery from the unoccluded shampoo dose group was high because the Coumarin was only applied for 5 minutes before being removed and recovered in the skin surface wash. The recovery of 4-HR after application in ethanol was only 70%; 4-HR isn’t considered to be volatile, so why this Mass Balance was so low is unclear, it might be that as the ethanol volatilised off, it took some 4-HR with it.

**Figure 9.**
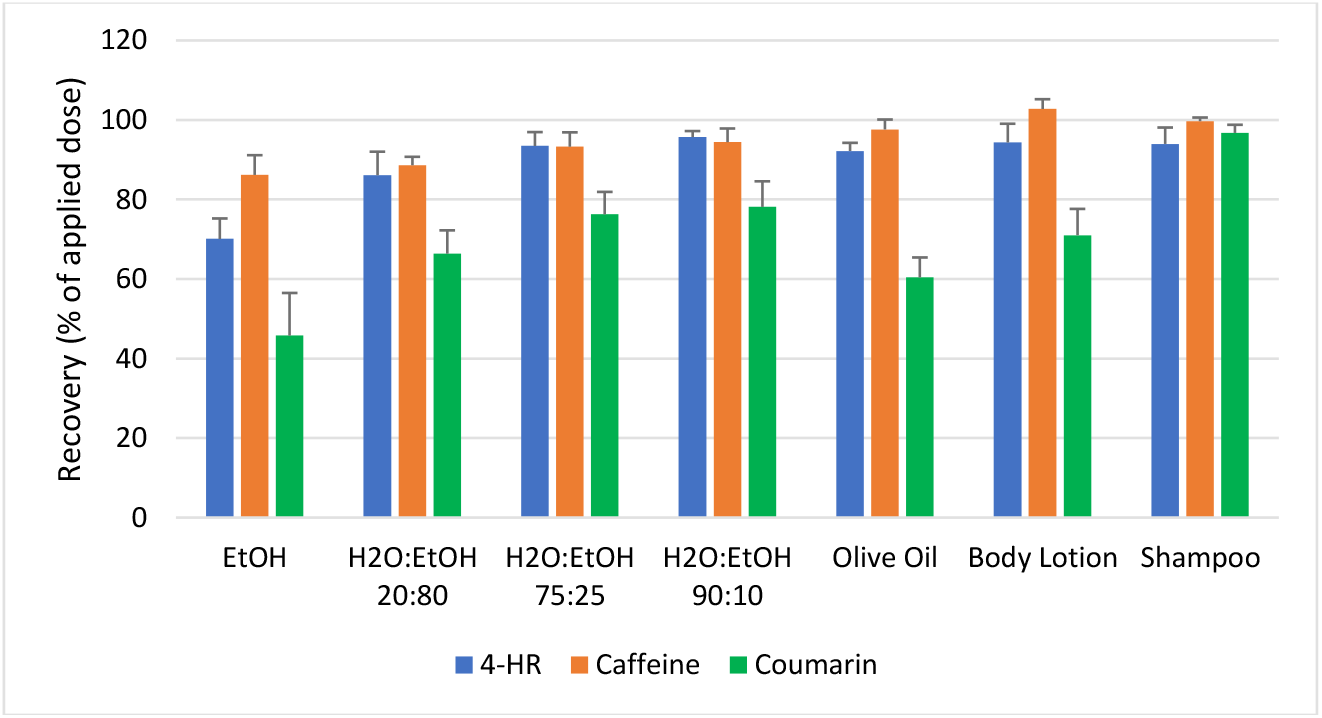
Mass balance for each dose group as a percentage of the applied dose

### Evaluation of the predicted skin penetration of the three chemicals in different vehicles with measured data

The values of diffusivity and partition coefficient parameters used in the TCAT model are presented in Table 5. The default value of vehicle water partition coefficient is 1, independent of which chemical is modelled. With the PDMS data (Table 4), a vehicle water partition coefficient could be calculated. Other parameters were obtained by fitting against the measured skin pen data in receptor fluid for all three chemicals when 10% ethanol was the vehicle. With these input parameters, skin penetration was modelled and compared with the measured dermal absorption data of 7 vehicle conditions (Figures 10-13). The vehicle water partition coefficient was set either at 1 or with the PDMS derived value for comparison.

**Table 5.** Diffusivity and partition coefficient values.

| Chemical | Vehicle | Diffusivity (cm/s<sup>2</sup>) <sup>a</sup> |  |  |  | Partition coefficient (vs water) <sup>a</sup> |  |  |  |  |  |
| --- | --- | --- | --- | --- | --- | --- | --- | --- | --- | --- | --- |
|  |  | Vehicle | Stratum corneum | Viable epidermis | Dermis | Vehicle |  | Stratum corneum | Viable epidermis | Dermis | Receptor fluid <sup>c</sup> |
|  |  |  |  |  |  | PDMS data <sup>b</sup> | default value |  |  |  |  |
| Caffeine | 10% EtOH | 1.2E-5 | 6E-11 | 2.2E-8 | 2.2E-6 | 1.3 | 1 | 0.7 | 0.7 | 0.7 | 0.7 |
|  | 25% EtOH |  |  |  |  | 2.4 |  |  |  |  |  |
|  | 80% EtOH |  |  |  |  | 8.6 |  |  |  |  |  |
|  | EtOH |  |  |  |  | 1.2 |  |  |  |  |  |
|  | Olive oil |  |  |  |  | 0.2 |  |  |  |  |  |
|  | Body lotion |  |  |  |  | 1.9 |  |  |  |  |  |
|  | SHAMPOO |  |  |  |  | 0.9 |  |  |  |  |  |
| Coumarin | 10% EtOH | 1.2E-5 | 4E-10 | 7E-8 | 3E-6 | 1.2 | 1 | 2 | 2 | 2 | 2 |
|  | 25% EtOH |  |  |  |  | 2.7 |  |  |  |  |  |
|  | 80% EtOH |  |  |  |  | 35.1 |  |  |  |  |  |
|  | EtOH |  |  |  |  | 21.3 |  |  |  |  |  |
|  | Olive oil |  |  |  |  | 11.3 |  |  |  |  |  |
|  | Body lotion |  |  |  |  | 1.9 |  |  |  |  |  |
|  | SHAMPOO |  |  |  |  | 1.0 |  |  |  |  |  |
| 4HR | 10% EtOH | 1.2E-6 | 5E-10 | 6E-10 | 1E-8 | 1.3 | 1 | 60 | 50 | 30 | 30 |
|  | 25% EtOH |  |  |  |  | 11.7 |  |  |  |  |  |
|  | 80% EtOH |  |  |  |  | 538.8 |  |  |  |  |  |
|  | EtOH |  |  |  |  | 144.2 |  |  |  |  |  |
|  | Olive oil |  |  |  |  | 182.6 |  |  |  |  |  |
|  | Body lotion |  |  |  |  | 135.7 |  |  |  |  |  |
|  | SHAMPOO |  |  |  |  | 39.2 |  |  |  |  |  |

<sup>a</sup> - values fitted against skin pen data of 10% ethanol (apart from the vehicle water partition coefficient which was derived from PDMS data or set at default value)
<sup>b</sup> - PDMS data calculated based on data in Table 4; <sup>c</sup> - receptor fluid partition coefficient set equal to that of dermis to assure it's not rate limiting

**Table 6.** Table of Errors in fits with 3 chemicals in tested formulations.

|  | 4HR |  | Caffeine |  | Coumarin |  |
| --- | --- | --- | --- | --- | --- | --- |
|  | RMS Error |  | RMS Error |  | RMS Error |  |
| **Formulation** | Original | PDMS | Original | PDMS | Original | PDMS |
| Ethanol | 8.7 | 1.6 | 31.5 | 27.2 | 15.8 | 9.2 |
| H2O EtOH 20/80 | 12.4 | 2.3 | 16.9 | 14.0 | 14.9 | 33.8 |
| H2O EtOH 75/25 | 5.3 | 4.0 | 8.2 | 6.8 | 3.5 | 11.9 |
| H2O EtOH 90/10 | 5.6 | 5.7 | 9.7 | 5.4 | 2.7 | 2.6 |
| Olive Oil | 17.8 | 2.9 | 15.4 | 55.8 | 51.3 | 5.8 |
| Shampoo | 2.4 | 0.1 | 0.0 | 0.1 | 1.2 | 1.2 |
| VICL | 14.2 | 1.7 | 4.9 | 5.6 | 14.3 | 8.5 |

**Figure 10.**
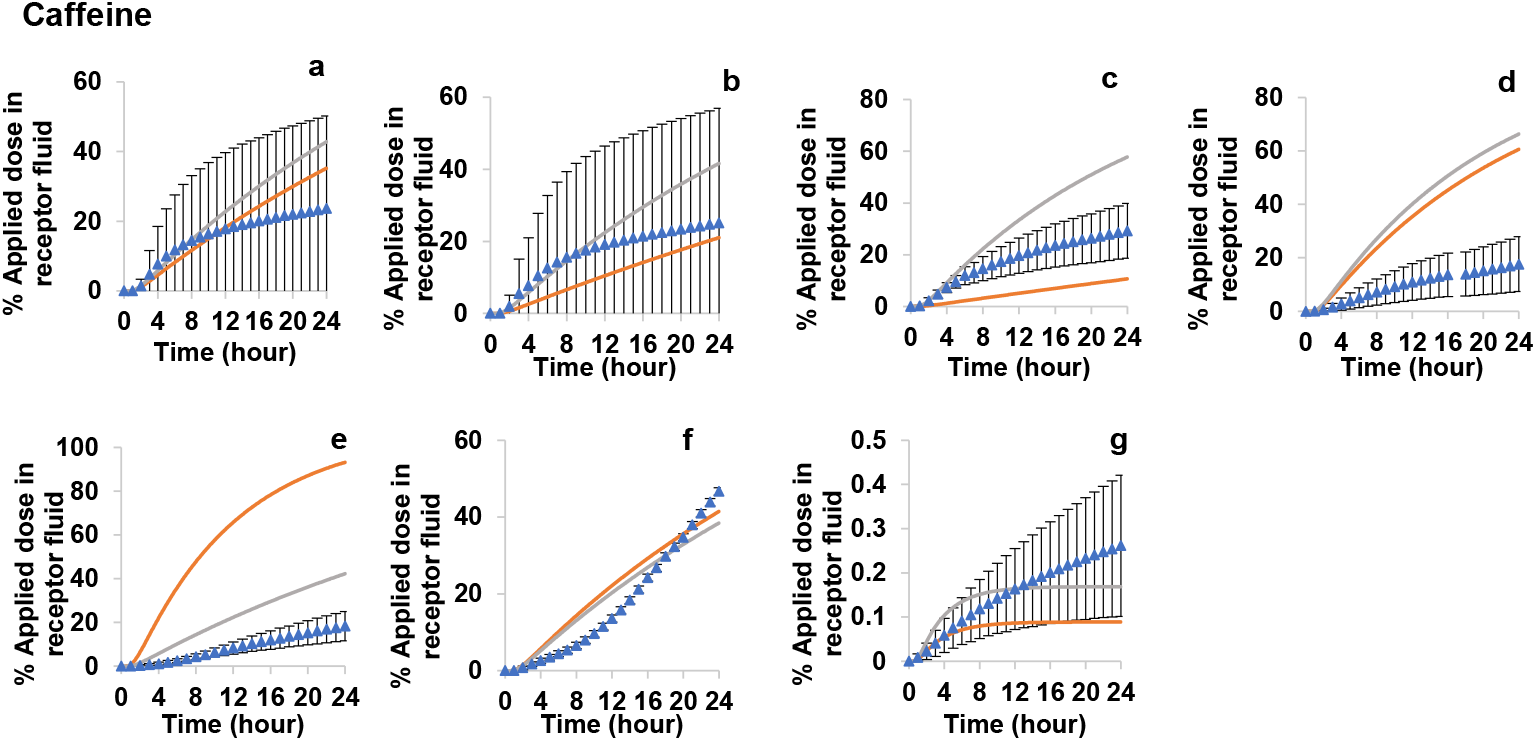
Caffeine modelling of dermal absorption data (% applied in receptor fluid over time) with different vehicles (a) 10% ethanol in water (which was used for fitting diffusivity and partition coefficient parameters in the skin layers), (b) 25% ethanol in water, (c) 80% ethanol in water, (d) 100% ethanol, (e) olive oil, (f) body lotion and (g)shampoo. Dots represent the measured data in dermal absorption studies, error bars represent standard deviation. Curves represent the prediction with (orange) and without (grey) PDMS data derived v/h partition coefficient.

**Figure 11.**
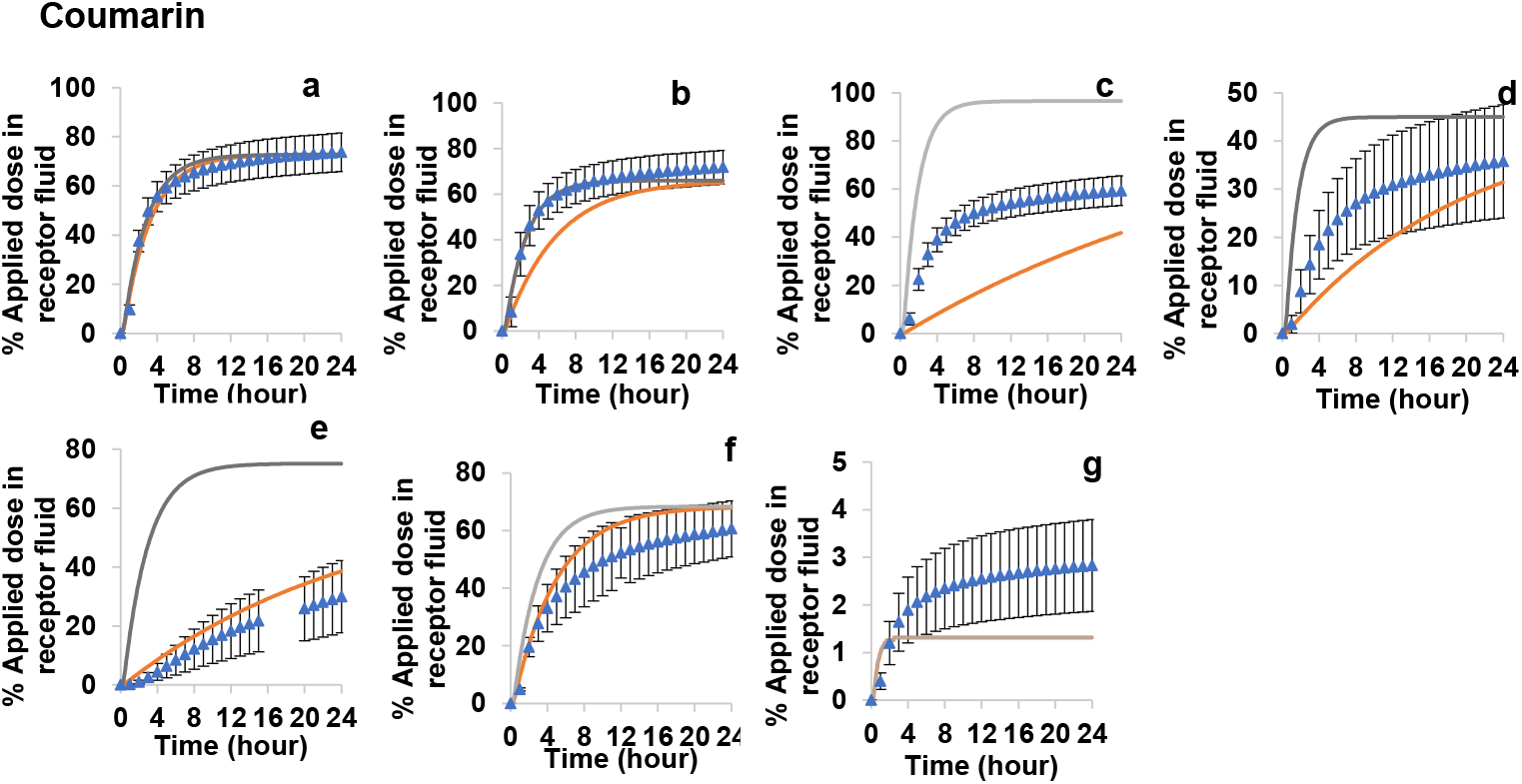
Coumarin modelling of dermal absorption data (% applied in receptor fluid over time) with different vehicles (a) 10% ethanol in water (which was used for fitting diffusivity and partition coefficient parameters in the skin layers), (b) 25% ethanol in water, (c) 80% ethanol in water, (d) 100% ethanol, (e) olive oil, (f) body lotion and (g)shampoo. Dots represent the measured data in dermal absorption studies, error bars represent standard deviation. Curves represent the prediction with (orange) and without (grey) PDMS data derived v/h partition coefficient.

**Figure 12.**
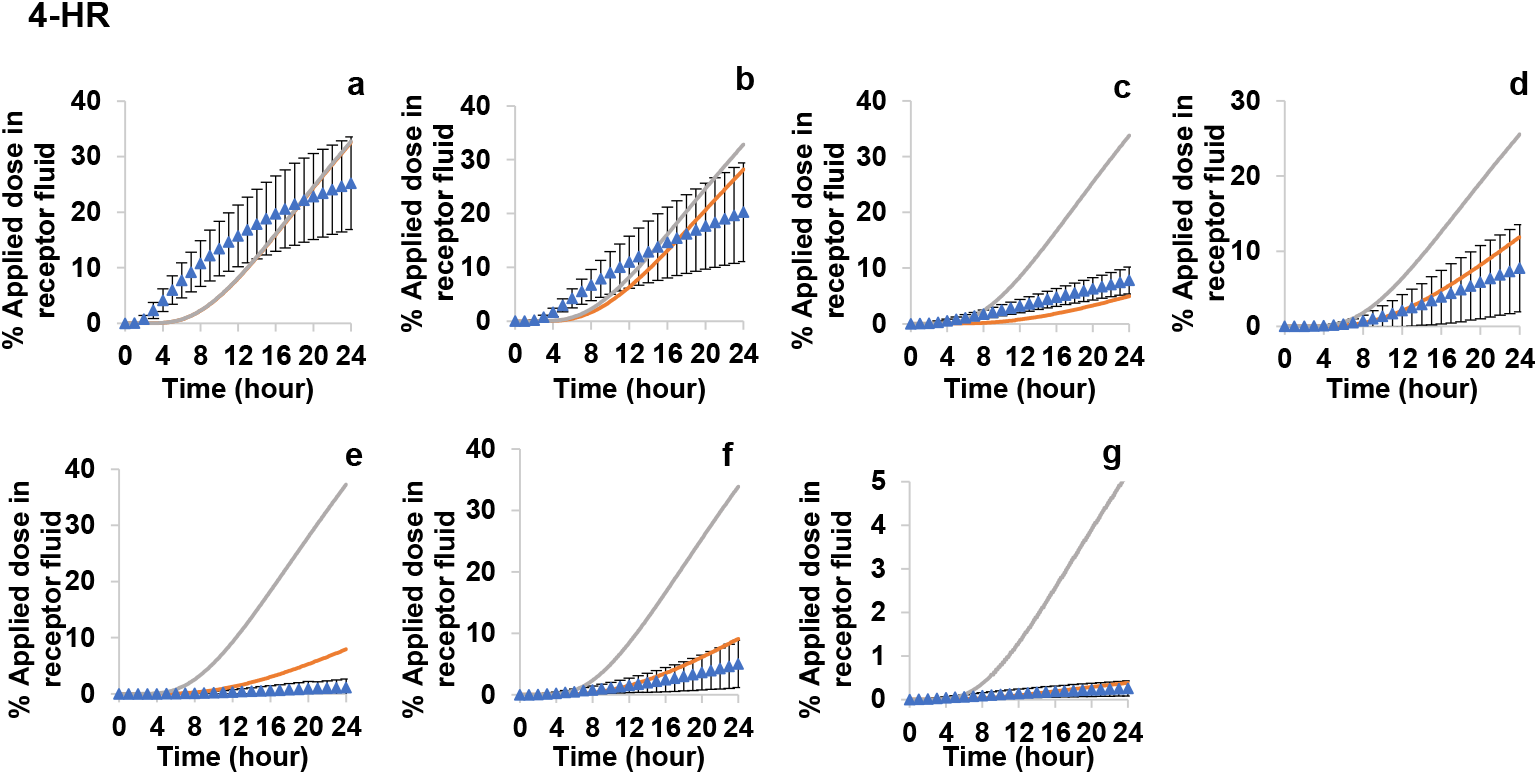
4-Hexylresorcinol modelling of dermal absorption data (% applied in receptor fluid over time) with different vehicles (a) 10% ethanol in water (which was used for fitting diffusivity and partition coefficient parameters in the skin layers), (b) 25% ethanol in water, (c) 80% ethanol in water, (d) 100% ethanol, (e) olive oil, (f) body lotion and (g)shampoo. Dots represent the measured data in dermal absorption studies, error bars represent standard deviation. Curves represent the prediction with (orange) and without (grey) PDMS data derived v/h partition coefficient.

**Figure 13.**
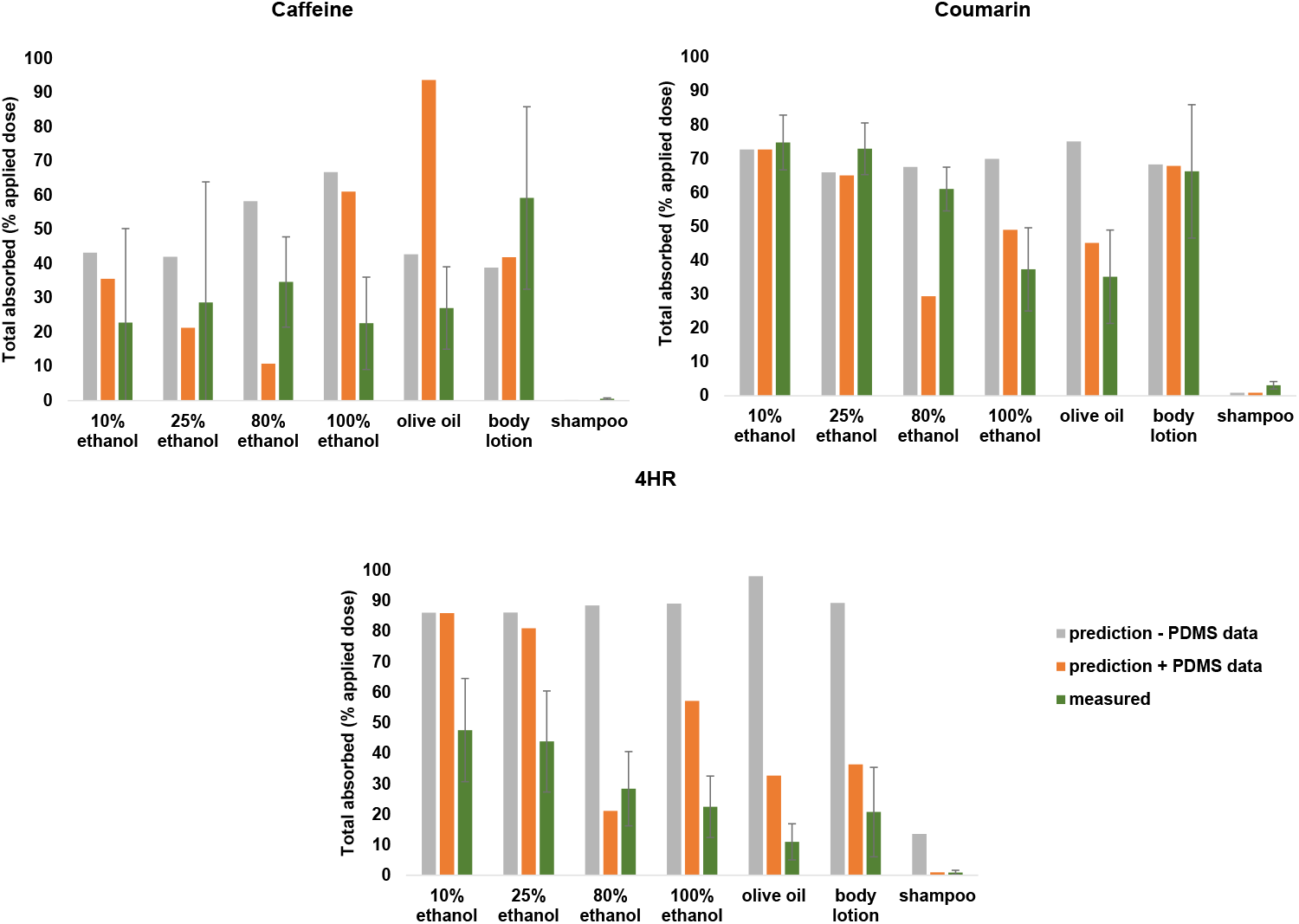
Comparison of predicted dermal absorption data (total absorption % applied at 24h) with (orange) and without (grey) PDMS data derived vehicle partition coefficient in different vehicles against measured data (green).

Figures 10-13 present the evaluation of the modelled skin penetration of the three chemicals using the TCAT module in different vehicles with measured skin absorption data. In general, PDMS data has shown to be promising in improving prediction on skin absorption for most of the conditions. For caffeine, with the PDMS derived parameter, the model showed to predict the percent recovered in receptor fluid better, except for the oil condition. For coumarin and 4-HR, the model with PDMS data has shown to improve the prediction on chemical absorption dramatically for all the vehicles except coumarin in 80% ethanol.

In Table 6 we show the errors in each fit before and after the use of the partition coefficient derived from the PDMS surrogate membrane. We see that for most test items and formulations a substantial improvement in the accuracy of the modelling is observed. For 4HR, demonstrated either a substantial increase in accuracy, or no substantial decrease as measured by the RMS error. In the case of caffeine we see a similar trend, bar in VICL and Olive Oil. Likewise in coumarin we observe that for most formulations, there is a an increase in the accuracy of the model predictions, except for the higher ethanol/water formulations.

## DISCUSSION

In nonclinical safety assessments of cosmetic ingredients applied topically, it is essential to investigate the rate and extent of dermal penetration to reach systemic circulation. Several factors can affect the absorption and distribution of substances in the skin, encompassing interactions between the vehicle and the skin, as well as interactions between the active ingredient and the skin. (Otto et al. 2009b). The initial step in skin absorption involves the partitioning of an ingredient from the formulation into the upper layer of the stratum corneum and plays a key role in determining the delivery of a chemical. Currently, there is a lack of reliable methods to accurately predict this important parameter. To address this, we measured the partitioning of an ingredient from seven different vehicles into a surrogate PDMS membrane. These measurements were then used to parameterize a dermal absorption model. Model predictions on skin absorption were then compared against our previously generated data from an ex vivo skin absorption study to evaluate the usefulness of the derived parameters from the PDMS method on formulation effect.

The three test items have a range of physicochemical properties (Table 3). Partitioning of the test items into the PDMS membrane was as anticipated considering the physicochemical properties of each one. For the more hydrophobic 4-HR, the partition coefficient between water and the PDMS membrane was high, indicating the 4HR preferentially partitions out of the water into the PDMS membrane, whilst for caffeine the PDMS water partition coefficient was much lower, with coumarin having an intermediate value (Table 4). At the other extreme, the PDMS/olive oil partition coefficient was very low for 4HR whilst for caffeine, this was the highest value measured for all the solvent systems, with coumarin again having an intermediate value.

The assessment of variability in the data sets showed that although only two vials were used per test item/vehicle combination for the partition coefficient measurement experiments, there was strong agreement between the two, and for those instances where there was a discrepancy between vials, a vial effect was fitted in the modelling phase to account for this. Furthermore, the results showed very small variance within test and material combinations compared to between test variance, meaning we can have high confidence in the robustness of the results.

The ability to accurately predict flux into the systemic circulation is important to predict an accurate plasma C_max_ value in PBK models, irrespective of whether the dose is applied orally or topically. In the topical situation, traditionally *ex vitro* skin absorption experiments have provided risk assessors with a total absorbed percentage value, whilst the kinetic flux data per se were often ignored, even though they are an important guide as to whether the conditions of exposure are finite or infinite. This is especially important in determining whether material in the stratum corneum at the end of the 24 hour experimental period should be regarded as bioavailable or not. If the system is at steady state at the 24 hour termination time (infinite dose), running the experiment for longer would result in more test item passing into the receptor solution; in this instance the stratum corneum acts as reservoir. Conversely, if there is a lot of material associated with the SC at the 24 hour termination time point, but the flux has reduced to baseline earlier in the experiment, the material in the stratum corneum is likely bound to it and will not be available for further permeation – this is usually the case for permanent hair dyes ingredients involved in the reactions that produce the dye’s colour; in this instance, material in the SC would be considered not available for absorption and would be excluded from the calculation of the absorbable dose.

The evaluation of receptor fluid kinetics and skin absorption percentages, comparing measured and predicted data, demonstrated that incorporating the vehicle partitioning parameter derived from the PDMS system greatly improved the accuracy of model predictions for three chemicals. However, two exceptions were identified: the predictions for caffeine in olive oil and coumarin in 80% ethanol exhibited a decline in accuracy when PDMS-derived data was included, resulting in inconsistencies with the measured data. Further investigations are necessary to ascertain the underlying cause of this discrepancy.

In this paper the other input parameter associated with the formulation that was considered was the evaporation of components from the vehicle. Evaporation of e.g. water from a formulation will increase the concentration of the remaining components on the skin surface, this could increase the permeation of an active by increasing its thermodynamic activity. Conversely, permeation could be decreased by evaporation of water by decreasing the partition coefficient between vehicle and stratum corneum if removal of water increases the solubility of the active in the residual formulation (Arce et al. 2020). In the modelling described in this paper, evaporation of vehicle components was addressed by making some assumptions using literature data and accounting for occlusion or non-occlusion of the skin absorption diffusion cells; measuring changes to a formulation in real time could give increased confidence to this input. The experimentally measured solubility of the test item in the specific vehicle (or relevant phase of the specific vehicle in the case of multiple phase vehicles) would be another more accurate input parameter for GastroPlus.

In general, PDMS data have shown to be promising in improving prediction on skin absorption for most of the conditions, however these conclusions are drawn for only three test items. The analysis of the overall accuracy of the fitting pre and post the PDMS correction suggests that although for most systems an improved accuracy was observed, there may exist some limitations of the method depending on the hydrophobicity of the test compound and relative hydrophobicity of the formulation in question. We note that for a hydrophilic compound in a lipophilic formulation, such as caffeine in olive oil, and for hydrophobic coumarin in ethanol/water the predictions of the model disimproved. However, for 4HR we saw an improvement for all formulations. Further tests with a wider range of chemicals and formulations may determine the physicochemical properties that determine how effective the PDMS surrogate method more fully.

## References

Arce F, Jr., Asano N, See GL, et al. (2020) Prediction of skin permeation and concentration of rhododendrol applied as finite dose from complex cosmetic vehicles. Int J Pharm 578:119186 doi:10.1016/j.ijpharm.2020.119186

Bernauer U, Bodin L, Chaudhry Q, et al. (2023) SCCS Notes of guidance for the testing of cosmetic ingredients and their safety evaluation - 12th revision - Final Opinion – SCCS/1647/22 - Corrigendum 2,

Chilcott RP, Barai N, Beezer AE, et al. (2005) Inter- and intralaboratory variation of in vitro diffusion cell measurements: an international multicenter study using quasi-standardized methods and materials. J Pharm Sci 94(3):632–8 doi:10.1002/jps.20229

Ellison CA, Tankersley KO, Obringer CM, et al. (2020) Partition coefficient and diffusion coefficient determinations of 50 compounds in human intact skin, isolated skin layers and isolated stratum corneum lipids. Toxicol In Vitro 69:104990 doi:10.1016/j.tiv.2020.104990

Grégoire S, Sorrell I, Lange D, et al. (2021) Cosmetics Europe evaluation of 6 in silico skin penetration models. Computational Toxicology 19 doi:10.1016/j.comtox.2021.100177

Hansen S, Henning A, Naegel A, et al. (2008) In-silico model of skin penetration based on experimentally determined input parameters. Part I: experimental determination of partition and diffusion coefficients. Eur J Pharm Biopharm 68(2):352–67 doi:10.1016/j.ejpb.2007.05.012

Li H, Reynolds J, Sorrell I, et al. (2022) PBK modelling of topical application and characterisation of the uncertainty of C(max) estimate: A case study approach. Toxicol Appl Pharmacol 442:115992 doi:10.1016/j.taap.2022.115992

OECD (2004) OECD 428 Guideline for the testing of chemicals. Skin Absorption: in vitro method.

Otto A, du Plessis J, Wiechers JW (2009a) Formulation effects of topical emulsions on transdermal and dermal delivery. Int J Cosmet Sci 31(1):1–19 doi:10.1111/j.1468-2494.2008.00467.x

Otto A, Du Plessis J, Wiechers JW (2009b) Formulation effects of topical emulsions on transdermal and dermal delivery. International Journal of Cosmetic Science 31(1):1–19 doi:10.1111/j.1468-2494.2008.00467.x

Pendlington RU, Minter HJ, Stupart L, et al. (2008) Development of a modified in vitro skin absorption method to study the epidermal/dermal disposition of a contact allergen in human skin. Cutan Ocul Toxicol 27(4):283–94 doi:10.1080/15569520802327005

Pendlington RU, Whittle E, Robinson JA, Howes D (2001) Fate of ethanol topically applied to skin. Food Chem Toxicol 39(2):169–74

Smith JM, Van Ness HC, Abbott MM, Swihart MT (2017) Introduction to Chemical Engineering Thermodynamics 8th Edition. McGraw-Hill Education

Supe S, Takudage P (2021) Methods for evaluating penetration of drug into the skin: A review. Skin Research and Technology 27(3):299–308 doi:10.1111/srt.12968

Van der Merwe D, Riviere JE (2005) Comparative studies on the effects of water, ethanol and water/ethanol mixtures on chemical partitioning into porcine stratum corneum and silastic membrane. Toxicol In Vitro 19(1):69–77 doi:10.1016/j.tiv.2004.06.002

Machatha SG, Yalkowsky SH (2005). Bilinear model for the prediction of drug solubility in ethanol/water mixtures. J Pharm Sci 94(12):2731–4. doi: 10.1002/jps.20492. PMID: 16258999.

OECD (2004) OECD 428 Guideline for the testing of chemicals. Skin Absorption: in vitro method.

